# Oligotyping and Genome-Resolved Metagenomics Reveal Distinct *Candidatus* Accumulibacter Communities in Full-Scale Side-Stream versus Conventional Enhanced Biological Phosphorus Removal (EBPR) Configurations

**DOI:** 10.1101/596692

**Authors:** Varun N. Srinivasan, Guangyu Li, Dongqi Wang, Nicholas B. Tooker, Zihan Dai, Annalisa Onnis-Hayden, Ameet Pinto, April Z. Gu

**Affiliations:** Department of Civil and Environmental Engineering, Northeastern University, Boston MA-02115, USA; State Key Laboratory of Eco-hydraulics in Northwest Arid Region, Xi’an University of Technology, Xi’an, Shaanxi 710048, China; Department of Civil and Environmental Engineering, University of Massachusetts-Amherst, Amherst MA-01002, USA; Infrastructure and Environment Division, University of Glasgow, Glasgow G12 8LT, United Kingdom; Civil and Environmental Engineering, Cornell University, Ithaca NY – 14853, USA

**Author notes:** **Corresponding Authors:** April Gu, Civil and Environmental Engineering, Cornell University.

## Abstract

*Candidatus* Accumulibacter phosphatis (CAP) and its sub-clades-level diversity has been associated and implicated in successful phosphorus removal performance in enhanced biological phosphorus removal (EBPR). Development of high-throughput untargeted methods to characterize clades of CAP in EBPR communities can enable a better understanding of Accumulibacter ecology at a higher-resolution beyond OTU-level in wastewater resource recovery facilities (WRRFs). In this study, for the first time, using integrated 16S rRNA gene sequencing, oligotyping and genome-resolved metagenomics, we were able to reveal clade-level differences in Accumulibacter communities and associate the differences with two different full-scale EBPR configurations. The results led to the identification and characterization of a distinct and dominant Accumulibacter oligotype - Oligotype 2 (belonging to Clade IIC) and its matching MAG (RC14) associated with side-stream EBPR configuration. We are also able to extract MAGs belonging to CAP clades IIB (RCAB4-2) and II (RC18) which did not have representative genomes before. This study demonstrates and validates the use of a high-throughput approach of oligotyping analysis of 16S rRNA gene sequences to elucidate CAP clade-level diversity. We also show the existence of a previously uncharacterized diversity of CAP clades in full-scale EBPR communities through extraction of MAGs, for the first time from full-scale facilities.

## Introduction

Enhanced biological phosphorus removal (EBPR) has been a promising technology for phosphorus removal from municipal wastewater, due to its sustainable nature and P recovery potential in comparison to chemical precipitation and adsorption-based approaches [1]. However, conventional EBPR processes face challenges related to system stability and susceptibility to variations and fluctuations in influent loading conditions such as unfavorable carbon to P (C/P) ratios [2]. A new emerging process, namely, side-stream EBPR (S2EBPR) features an anaerobic side-stream reactor that allows a portion of the return activated sludge (RAS) to undergo hydrolysis and fermentation thus enabling influent carbon-independent enrichment and selection of polyphosphate accumulation organisms (PAO). Facilities that have operated or piloted S2EBPR showed improved and more stable performance [3–6]. However, several key knowledge gaps still exist for this process including lack of understanding of the fundamental mechanisms governing the process and differences in microbial ecology, particularly functionally relevant key populations such as PAOs between the conventional and the S2EBPR processes. Several studies that have characterized the microbial ecology of different process configurations using 16S rRNA gene amplicon sequencing, fluorescence in-situ hybridization (FISH) and quantitative polymerase chain reaction (qPCR) have failed to associate differences in PAO microbial ecology with process configurations [7–10]. This is, at least partially due to the challenges in identification and association of specific PAO populations with various full-scale EBPR configurations. The complex and overwhelming “masking” effect of facility-specific selection forces exerted on the microbial community structure makes it difficult to isolate the effect of process configuration alone.

*Candidatus* (*Ca*.) Accumulibacter phosphatis (hereafter referred to as Accumulibacter or CAP) is considered to be a key PAO in both lab-scale and full-scale EBPR systems [11–16]. Our current understanding of the physiology, metabolic potential, transcriptional, and proteomic dynamics are based on lab-scale Accumulibacter-enriched reactors from which 19 metagenome-assembled genomes (MAGs) of Accumulibacter have been obtained [12, 16–20] and amongst them, only one complete draft genome sequence for Clade IIA, namely Accumulibacter UW1 has been reported. These MAGs have highlighted the enormous genomic diversity in Accumulibacter clades (>700 unique genes in each clade) especially in terms of the capacity to perform the different steps of the denitrification pathway and ability to metabolize complex carbon sources [16]. However, besides the MAGs from lab-scale reactors, Accumulibacter genome assemblies from full-scale systems have not been reported, possibly due to their low relative abundance (< 5 % based on 16S rRNA gene sequencing) [10, 14] and insufficient depth of sequencing, which makes MAG extraction from metagenome assemblies difficult. Metagenomic studies have indicated high-level of diversity within Accumulibacter populations in full-scale EBPR communities [10, 14], however this diversity was not fully characterized due to the inability to extract high-quality MAGs. This phylogenetic diversity within Accumulibacter populations along with the genomic functional diversity across clades points towards ecological niche-partitioning due to determinants such as type of carbon source, phosphorus levels, length of anaerobic period, and presence of alternate electron acceptors such as nitrate and nitrite etc. Characterization of the microdiversity along with generation of corresponding MAGs could lead to a better understanding of the niche determinants of Accumulibacter ecology which would help better inform treatment process optimization and operation.

Current methods to characterize clade-level differences in Accumulibacter populations include qPCR, FISH, *ppk1* gene sequencing and metagenomic sequencing. qPCR and FISH have been extensively used to characterize the clade-level differences in lab-scale and full-scale systems [21–24]. However, they target only those currently known clades for which primers or probes are available. Metagenomic and *ppk1* gene sequencing are untargeted analyses and are capable of providing phylogenetic information, however they are cost-prohibitive and/or time-consuming for routine analysis at full-scale practice. Although, 16S rRNA gene amplicon sequencing has emerged to be a high-throughput and widely-used method to characterize and capture broad shifts in microbial communities, it is recognized that it fails to capture the phylogenetic diversity within OTUs (also called microdiversity) [25, 26]. Recent studies have highlighted the importance of capturing this diversity and that genetic variation in 16S rRNA gene amplicon sequences can capture meaningful ecologically distinct units [27, 28]. Recent work has argued for the use of a higher similarity threshold [25, 26] or for the use of exact sequence variants (ESVs) or amplicon sequence variants (ASVs) [29, 30] to more accurately characterize the microdiversity in a community. Oligotyping has been proposed as a method that can distinguish meaningful sequence variation from sequencing errors and can partition sequence types that differ in as little as one nucleotide with the assumption that the variation in small regions of the 16S rRNA gene translate to phylogenetic differences [27, 28, 30].

In this study, we employed 16S rRNA gene sequencing, oligotyping and metagenomic analysis to elucidate and compare the microbial ecology and potential functional traits of Accumulibacter between conventional and S2EBPR process configurations in full-scale systems. The unique opportunity to operate two different EBPR configurations, in parallel and separate treatment trains with the same influent feed, made it possible to eliminate other influent characteristics-related selection factors and, therefore enabled association of specific CAP clades with S2EBPR versus conventional A2O configuration. To overcome the limitation of 16S rRNA gene sequencing and OTU-based approaches in revealing microdiversity, we demonstrated and validated the use of oligotyping in revealing the microdiversity within Accumulibacter populations. The results of the oligotyping analysis were further compared and confirmed with phylogenetic analysis of both full-length 16S rRNA and *ppk1* gene sequences retrieved from metagenomic sequencing. In addition, using differential coverage binning, we successfully assembled and binned 2 high-quality and 1 medium-quality [31] Accumulibacter MAGs from full-scale EBPR sludge for the first time, including a distinct *Ca*. Accumulibacter clade that is dominant and associated with the S2EBPR configuration. Lastly, comparative genomic analysis was performed with the extracted MAGs and with previously published MAGs, to determine differences in key metabolic pathways and functions including unique functions that could contribute to niche-partitioning in the dynamic and complex environments in full-scale EBPR facilities.

## Materials and Methods

### Rock Creek Pilot

Full-scale testing was conducted from June 22^nd^ to August 30^th^ 2016 at the Rock Creek Facility (Hillsboro, Oregon). Two parallel trains with separated secondary clarifiers were operated with the same influent wastewater side-by-side, with one operated as conventional EBPR and another one as S2EBPR configuration for comparison. The S2EBPR configuration implemented was side-stream RAS fermentation with supplemental carbon addition (SSRC). In this configuration, 100% of the RAS was diverted to the side-stream reactor (Figure S1) for a total anaerobic HRT of 1.5 hours with primary sludge fermentate addition. The side-stream reactor was mixed intermittently once every week for 10 minutes during the sampling period for this study. The conventional EBPR configuration was an A2O process with an anaerobic HRT of 0.7 hours. Average influent flow to each treatment trains during the testing period was 5.35 MGD. Throughout the testing period, process performance parameters were routinely monitored. Detailed information on operation and performance is available in supplementary methods and published literature [4, 6, 32].

### Sampling, DNA Extraction, Amplicon Sequencing and Analysis

Sampling was performed before pilot testing was started as a baseline (April 7th and April 26^th^, 2016) and during pilot testing (June 22^nd^, July 20^th^, August 2^nd^, August 17th, and August 30^th^, 2016). MLSS samples were collected at the end of the aerobic zone, shipped to Northeastern University, Boston MA overnight on dry ice. DNA was extracted according to the MiDAS protocol [33]. The extracted DNA was sent to University of Connecticut-MARS facility for PCR amplification and sequencing targeting the V4 region using the primers 515F (5’-GTGCCAGCMGCCGCGGTAA-3’) and 806R (5’-GGACTACHVGGGTWTCTAAT-3’) [34] and the amplicons were sequenced on the Illumina MiSeq using V2 chemistry using paired-end (2 × 250) sequencing [35]. Raw reads have been submitted to NCBI under the BioProject accession number PRJNA530271.

The sequences were trimmed to remove primers and barcodes, quality filtered using sickle v1.33 [36] with a minimum quality score of 20 and analyzed as described in Kozich et al., [35]. Consensus taxonomy of OTUs was determined using the 80% cugtoff using the MiDAS (v123) database [33] (details of the analysis are provided in the supplementary methods). Amplicon data was rarefied to the minimum total sequence count (12090 sequences) across all samples. All data and statistical analysis were performed in R (for more details see supplementary methods) using the following packages: vegan (Oksanen et al. 2007), ggplot2 [37], dplyr [38] and ampvis [39].

### Oligotyping Analysis

To generate the input sequences required for oligotyping analysis, the Kozich et al., [35] pipeline, described above, was executed without the preclustering step as recommended by Eren et al. [30]. After chimera removal and denoising sequences, the 16S rRNA gene amplicon (V4 region) sequences classified as *Ca*. Accumulibacter were extracted and formatted as required by the oligotyping pipeline using the mothur2oligo.sh script (https://github.com/michberr/MicrobeMiseq/tree/master/mothur2oligo). A total of 38855 sequences that were classified as *Ca*. Accumulibacter were extracted and used as input for oligotyping analysis. The oligotyping pipeline (v2.1) was used according to Eren et al. [30] with recommended best practices. Quality filtering in the oligotyping pipeline resulted in 37934 reads (97.6 %). The Shannon entropy at each nucleotide position was calculated using the entropy-analysis command. Starting with the highest entropy positions, 9 nucleotide positions (56, 57, 63, 75, 112, 113, 114, 115, 123) were selected for entropy decomposition (Figure S2). Noise filtering was performed by limiting our analysis to oligotypes that occurred in at least 3 samples (-s) and those that had a minimum count (-M) of 30 (∼0.1 %). A total of 195 raw oligotypes were initially identified based on the 9 nucleotide positions. Denoising and elimination based on the -s and -M parameters resulted in 9 oligotypes.

### Metagenomic Sequencing, Assembly and Genome Binning

Samples from June 26^th^, August 2^nd^ and August 30^th^, 2016 from both treatment trains were sent to the TUCF Genomics center for 2 × 250 paired end sequencing on the Illumina HiSeq 2500. Library preparation for metagenomic sequencing was performed using the TruSeq PCR-free DNA kit. A total of 83.2 million paired-end reads were obtained. Sequences were quality filtered using sickle v1.33 [36] with a quality threshold of 20 and a length threshold of 50. In order to remove contamination sequences, the quality filtered sequences were then mapped to the UniVec database [40] and mapped reads were removed. Paired-end reads for all 6 samples were then co-assembled together using MEGAHIT [41] using a minimum contig length of 1000 to enable recovery of MAGs and eliminate erroneous contigs. Samples (n=3) from each treatment train were also co-assembled separately. Quality for the assemblies (total 3 assemblies) was assessed using QUAST [42]. The paired-end reads obtained after Univec-filtering were mapped to the assemblies using Bowtie 2.0 [43]. The resulting sam files were converted to bam, sorted and indexed using samtools [44]. Duplicate reads were removed using Picard [45]. Assembled contigs were then classified using Kaiju [46] with the default database. Genome binning was performed using the CONCOCT [47] implementation in Anvi’o [48] with a cluster size of 350. Resulting bins with >70 % completion were then manually refined in Anvi’o based on coverage patterns, GC content and taxonomic affiliation. The quality of all refined bins was then assessed with CheckM v1.0.11 and taxonomic classification was also obtained. Genome bins from the different assemblies were dereplicated using dRep [49] and the dereplicated bins were then used for downstream analysis. Bins, classified as *Rhodocyclaceae*, were extracted and paired-end reads from each sample were mapped against the MAGs using bbmap (v37.78) with 90% minimum identity while choosing only the best hit and matching paired-end reads. Contig-wise coverage was calculated using the pileup.sh script (https://github.com/BioInfoTools/BBMap/blob/master/sh/pileup.sh). Relative abundance of genome bins was calculated using the following equation (assuming an average genome length of 6 Mbp):

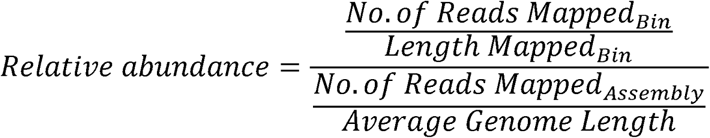

Raw reads and identified Accumulibacter MAG assemblies have been submitted to NCBI and are accessible under the BioProject accession number PRJNA530189.

### Reconstruction of full-length 16S rRNA Gene Sequences

EMIRGE [50] was used to reconstruct 16S rRNA gene sequences from the paired-end metagenomic data. Exact sequence matches to oligotypes were extracted, aligned with the reference sequences from He et al., [21] and a phylogenetic analysis was performed using MrBayes 3.2 [51] (see supplementary methods for details).

### ppk1 Annotation and Phylogenetic Analysiss

*ppk1* genes were annotated in the assemblies using a HMM model [52] using HMMER 3.1b2 (http://hmmer.org/). The annotated genes were then aligned with the reference sequences (n=1012) using MUSCLE v3.8.31 [53] and placed on a reference backbone tree using pplacer [54]. Read counts for each of the identified ppk1 sequences were obtained using the multicov tool in BEDtools [55] and the reads per kilobase million (RPKM) value was calculated. Sequences with length less than 600 bp were filtered out and relative abundance of the *ppk1* gene sequences was calculated using the following equation:

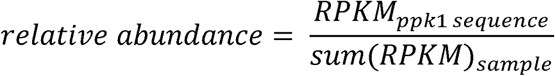

### Pangenomic and Phylogenetic Analysis

All publicly available genomes for *Ca*. Accumulibacter (n=19) were downloaded and combined with the CAP MAGs obtained in this study to perform a pangenomic analysis using the Anvi’o 5.1 [48] pangenomic workflow. Functional annotation of the MAGs was performed using GhostKOALA [56]. A set of 37 marker genes were identified using Phylosift and a concatenated alignment of these were used to construct a maximum-likelihood (ML) tree in RaxML version 8.2.11 [57] with a 100 bootstraps and with GTRGAMMA settings. For the phylogenetic analysis, published *Propionivibrioeu* (*Propionivibrio dixarboxylicus, Propionivibrio militaris, Ca*. Propionivibrio aalborgensis) and *Dechloromonas aromatica* RCB genomes were used as outgroups. The best-scoring ML tree was used. The *ppk1* gene was also annotated in all the genomes and used for a phylogenetic analysis using pplacer as outlined previously.

## Results and Discussion

### No observed differences in Accumulibacter diversity and relative abundance were observed at genus or OTU-level between conventional A2O and S2EBPR SSRC configurations

In this study, Accumulibacter was found to be the most abundant PAO in both EBPR configurations (Figure S2) [4, 6]. The estimated relative abundance of Accumulibacter, based on 16S rRNA gene sequencing, increased from the start of the pilot (June 21^st^, 2016) until the end (August 30^th^, 2016) in both treatment trains from an average abundance of 0.35 ± 0.05 % (both conventional and S2EBPR) to 1.98 ± 0.12 % (both conventional and S2EBPR) [4, 6]. A genus-level analysis did not reveal significant differences between the two EBPR configurations in the diversity and relative abundance of putative PAOs such as Accumulibacter. At the OTU-level, both treatment trains showed a predominance of OTU 00015 with most of the total Accumulibacter relative abundance accounted for by OTU 00015 (Figure 1). The rest of the OTUs were present at very low relative abundances. The results suggested that the diversity in Accumulibacter populations, if any, could not be resolved using OTU-based methods. However, there is still potential for finer-resolution diversity which could be masked within the OTUs.

**Figure 1:**
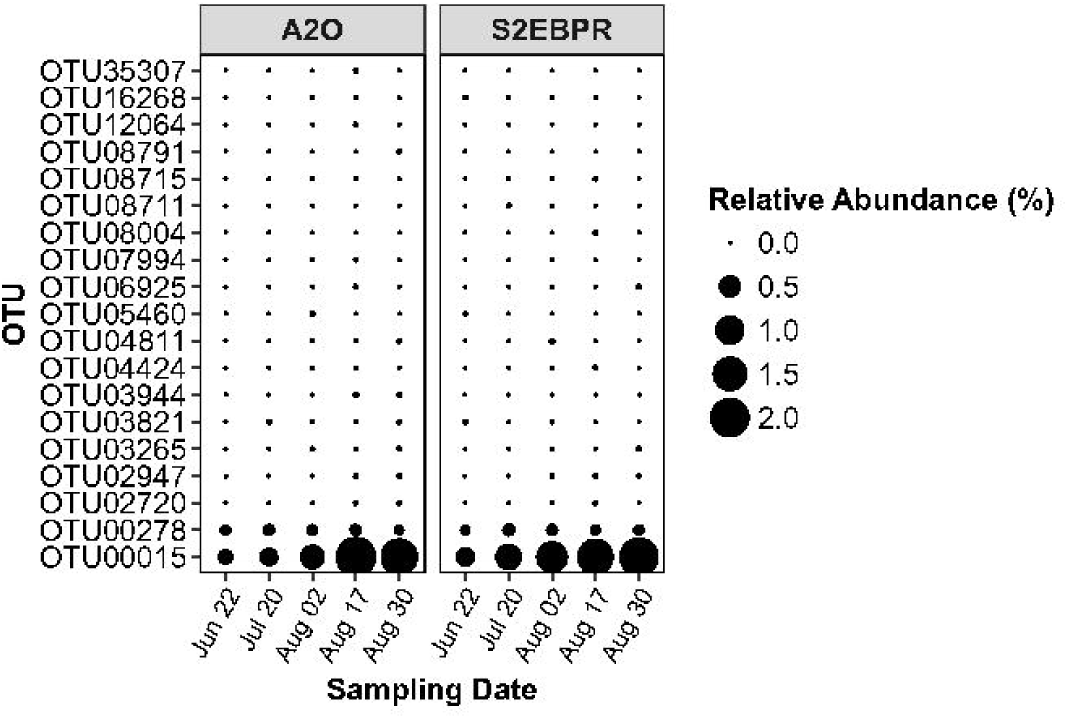
Comparison of most abundant OTUs classified as *Ca*. Accumulibacter in Conventional A2O and S2EBPR SSRC configurations.

### Oligotyping reveals differences in Accumulibacter clades between S2EBPR SSRC and conventional A2O configurations

In order to further resolve the microdiversity within Accumulibacter OTUs, oligotyping analysis was performed using the individual 16S rRNA gene sequences targeting the V4 region. The oligotyping analysis enabled the identification of 9 oligotypes of Accumulibacter and therefore distinct Accumulibacter community structure between the two EBPR configurations. The S2EBPR Accumulibacter community became increasingly dominated by Oligotype 2, during the testing period, while it remained at a very low relative abundance in the conventional A2O configuration (Figure 2A, Table S1). The relative abundance of Oligotype 2 increased from 31.8 ± 8.1 % at the start of testing (Jun 22) to 68.0 ± 0.2 % on Aug 17th and 62.4 ± 1.3 % on Aug 30th in S2EBPR while it remained below 20 % of the Accumulibacter sequences in the parallel conventional A2O treatment trains. The continuously increasing abundance of the Oligotype 2 in S2EBPR over a time-series and its consistently low abundance in A2O suggests that this oligotype was most likely favored and enriched by conditions in the S2EBPR configuration. In comparison, predominant oligotypes in A2O were Oligotypes 5 and 6 (Figure 2A). A hierarchical clustering analysis of the samples based on the oligotyping data showed distinct clusters based on operating regime (Figure S4). All samples from the S2EBPR train clustered together except for the sample from June 22nd which was the date when the piloting was started. Samples from A2O taken during the later stages of piloting (Aug 17th, Aug 30th) clustered together while the ones during the earlier stages (Jun 22nd, Jul 20th, and Aug 2nd) clustered separately.

**Figure 2:**
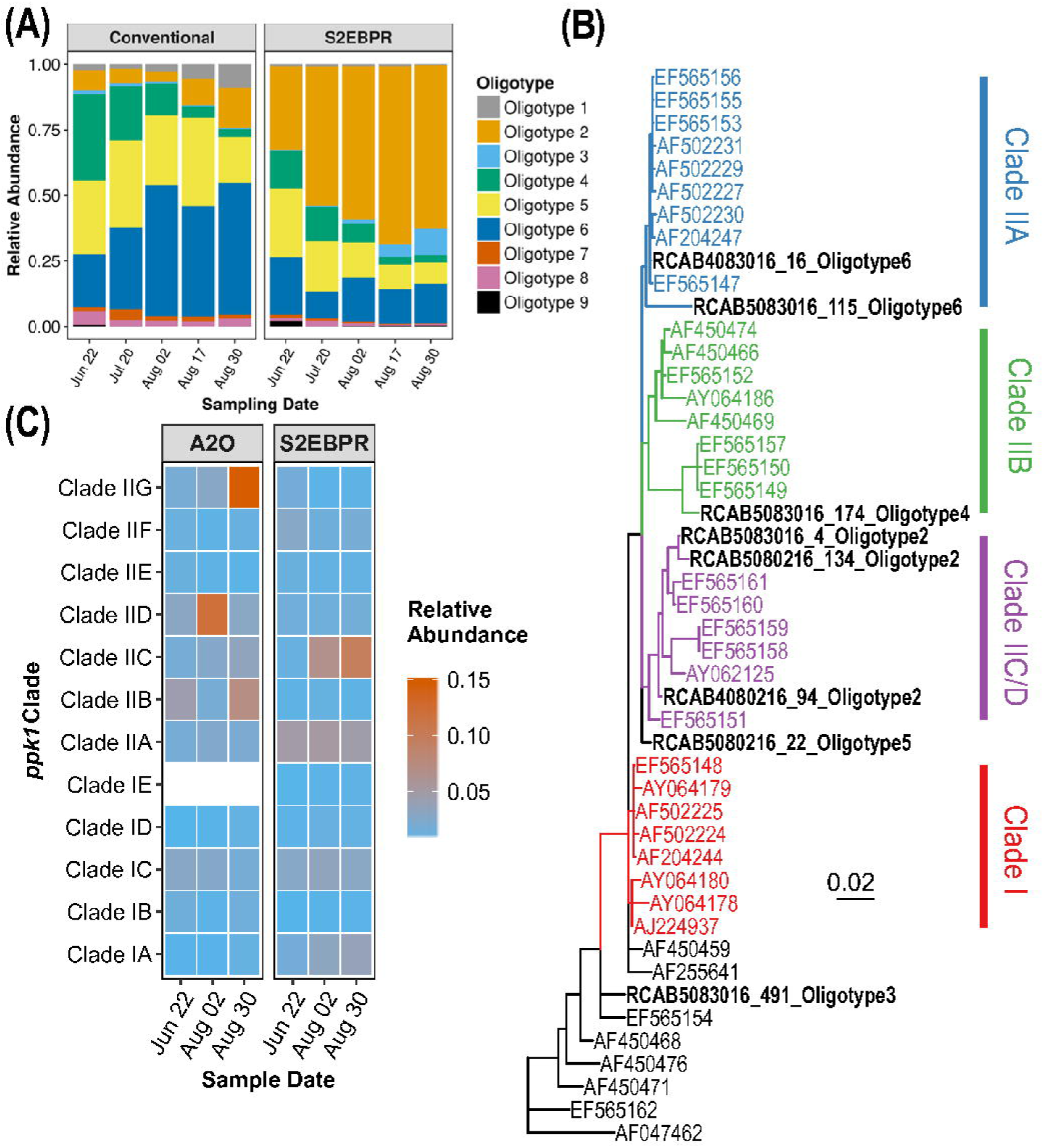
(A) Relative abundance of oligotypes of 16S rRNA gene amplicon sequences classified as *Ca*. Accumulibacter in conventional A2O and S2EBPR SSRC configurations (B) Phylogenetic tree of near full-length 16S rRNA gene sequences, that are exact matches to oligotypes, derived from metagenomic sequences using EMIRGE (highlighted in bold) and reference 16S rRNA gene sequences from He et al., (2007). Clade assignments for reference sequences are according to He et al., (2007) which were based on congruency with *ppk1* phylogeny (C) Relative Abundance of *ppk1* gene sequences extracted from the individual treatment-train assemblies and assigned clade-classification using pplacer.

### Phylogenetic analysis of oligotype sequences shows congruency with ppk1-phylogeny of Accumulibacter

In order to further correlate the oligotypes to clade/species level differences in CAP and examine the congruency of the oligotype phylogeny with *ppk1* phylogeny, we performed phylogenetic analysis using near full-length 16S rRNA gene and *ppk1* sequences through metagenomic sequencing on the samples taken on June 22nd, August 2nd, and August 30th from both treatment trains. A phylogenetic analysis of near full-length 16S rRNA gene sequences (extracted from metagenomic sequencing), classified as Accumulibacter and matching 5 of 9 oligotypes, along with reference sequences from He et al., [21], which were previously assigned to clades based on congruency with *ppk1* phylogeny, revealed that the 16S rRNA gene sequences matching the oligotypes were monophyletic (Figure 2B). It is important to note that multiple full-length 16S rRNA gene sequences with identical V4 regions matching each oligotype were obtained and they all clustered within the same clade. The sequences matching Oligotype 2, which is the dominant oligotype in S2EBPR, clustered with Clade IIC/D 16S rRNA gene sequences (He et al., 2007). Oligotype 6 clustered with Clade Clade IIA 16S rRNA gene sequences. Oligotype 5 did not cluster with any representative 16S rRNA reference sequence at a sub-clade level, but clustered with Clade II sequences. This could be due to the lack of a representative reference sequences for that particular sub-clade.

Sub-clades within Accumulibacter has been previously resolved using the *ppk1* gene [17, 58–61]. A phylogenetic analysis of *ppk1* gene sequences, extracted from metagenomic sequencing and assigned clades using pplacer (Figure S5), was performed and the cumulative relative abundance of sequences belonging to each clade was calculated (Figure 2C). This further confirmed that Clade IIC was predominant in S2EBPR which is congruent with oligotyping results. However, according to *ppk1* phylogeny, Clade IIG was the predominant clade in the conventional A2O configuration at the end of testing which was not directly congruent with the results using 16S rRNA oligotyping phylogeny. This could be due to the lack of representative 16S rRNA gene reference sequences for Clade IIG.

Previous research has cautioned that exact sequence variants (ESVs) or oligotypes may not represent phylogenetically or ecologically significant units [62]. However, this was based on a study of *Microcystis* in freshwater systems where they observed that the oligotypes observed in the study were not monophyletic and did not correlate with toxin producing capability. In this study, we show that the dominant oligotypes observed are distinct between EBPR configurations fed with the same influent wastewater and their full-length 16S rRNA gene sequence matches from metagenomic reconstruction are monophyletic. Furthermore, we show that the oligotyping results agree with the phylogenetic analysis with a more robust marker gene (*ppk1*) for Accumulibacter. It should be noted that, until now, the only way to resolve clade-level differences in Accumulibacter has been using the *ppk1* gene or using metagenomics. Even though qPCR primers for the *ppk1* gene exist for many clades, it is unclear whether these capture the diversity of Accumulibacter sufficiently. The potential advantages of resolving clade-level differences using a high-throughput method such as 16S rRNA gene amplicon sequencing combined with oligotyping is significant since it will enable more cost-effective and improved characterization of EBPR communities. However, the use of this method needs to be further validated by comparing with other tools that can differentiate closely-related taxa in order to characterize the microdiversity in a particular taxon.

### Accumulibacter MAGs binned from the metagenomic sequences reveal previously uncharacterized diversity in full-scale EBPR systems

To further characterize the identity and metabolic capabilities of Accumulibacter present in the two full-scale EBPR systems with different configurations, we used genome-resolved metagenomics to recover 3 MAGs (RC14, RC18, and RCAB4-2) classified putatively as Accumulibacter. The taxonomic affiliation of these MAGs was further delineated by performing a phylogenomic analysis along with 19 published Accumulibacter MAGs [13, 17, 18, 20, 63] using a concatenated alignment of 37 single-copy genes (SCG). Based on the SCG phylogeny (Figure s3A), the three MAGs were confirmed to be affiliated with Accumulibacter. RC14 and RC18 were high-quality MAGs with a completion of 99 % and 98.2 % respectively with low redundancy values of 0.2 % and 0.0 % respectively. The other MAG (RCAB4-2) had a completion of 77.2% and 3.1% redundancy.

To further delineate the clade-level taxonomy of the MAGs, the *ppk1* gene sequence from each MAG was extracted and placed on the reference tree as described previously. Based on their placement (Figure s3B), MAG RC14 can be classified as Clade IIC and RCAB4-2 as Clade IIB. Based on both SCG and *ppk1* phylogeny, MAG RC18 does not cluster under any of the canonical groupings at the sub-clade level, which suggests that it is a previously uncharacterized sub-clade of Accumulibacter since MAGs from full-scale facilities have not been published before. Both the SCG and *ppk1* phylogeny, suggest that RC18 would belong to Clade II. Average nucleotide identity (ANI) was used to confirm the results of the phylogenetic analysis (Figure S7). MAG RC14 had a > 90% ANI with Clade IIC MAGs over an alignment fraction of 57.0 % (BA91), 75.3 % (HKU2), 62 % (SK01), 70.8 % (SK02) and 72 % (UBA5574). This would confirm the classification of RC14 as a Clade IIC genome [64]. MAG RC18 and RCAB4-2 had an alignment fraction of < 10 % and < 20 % respectively with any of the other MAGs with ANIs of < 85 % again reinforcing the results of the phylogenetic analysis that these are previously uncharacterized clades [64]. We also calculated the relative abundance of these MAGs in the metagenome and observed that RC14 (Clade IIC) was the predominant Accumulibacter MAG in the S2EBPR metagenome (Figure 3C). Note that the clade affiliation of these genome bins corresponds to the predominant clades observed using both oligotyping and *ppk1* annotation (Figure 2). Clade IIC MAG (RC14) was the most abundant Accumulibacter MAG in S2EBPR, thus, again, confirming the oligotyping approach for resolving the microdiversity within CAP phylogeny.

**Figure 3:**
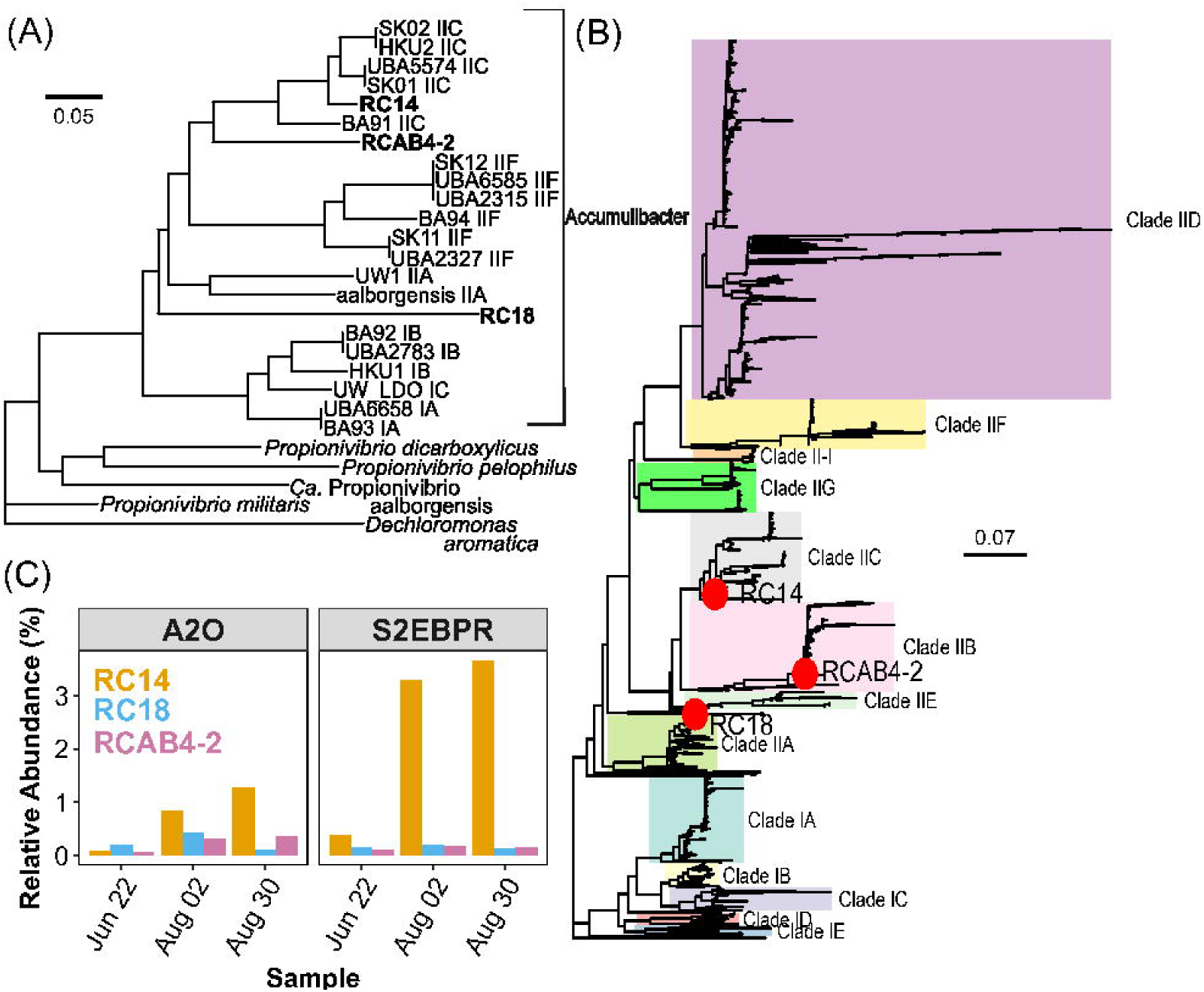
(A)Phylogeny of Ca. Accumulibcter MAGs recovered from this study along with published draft genomes (n=19) of Accumulibacter, *Propionivibrio* and *Dechloromonas aromatica* as an outgroup based on a concatenated alignment of 37 marker genes. Genome bins recovered in this study are in bold. (B) *ppk1* genes from Ca. Accumulibacter genomes recovered from this study placed on a backbone reference tree created with 1012 sequences and classified according to He et al.,[21] (C) Relative abundance of MAGs recovered in this study.

The newly identified RC18 genome seems to belong to a previously uncharacterized clade while RCAB4-2 is classified as Clade IIB for which there have not been any representative MAGs published. Our results revealed and highlighted the as-yet uncharacterized diversity of Accumulibacter. Particularly, draft genomes from full-scale systems have previously not been published, therefore the MAGs obtained in this study could expand the current knowledge on the genomic diversity in Accumulibacter in full-scale EBPR systems.

### Comparative Genomics of Ca. Accumulibacter

A comparative genomic analysis of 3 CAP MAGs obtained in this study along with 19 published CAP MAGs was performed with a focus on pathways relevant to EBPR metabolism such as polyP metabolism, phosphate transporters, carbon cycling, and denitrification (Figure 4).

**Figure 4:**
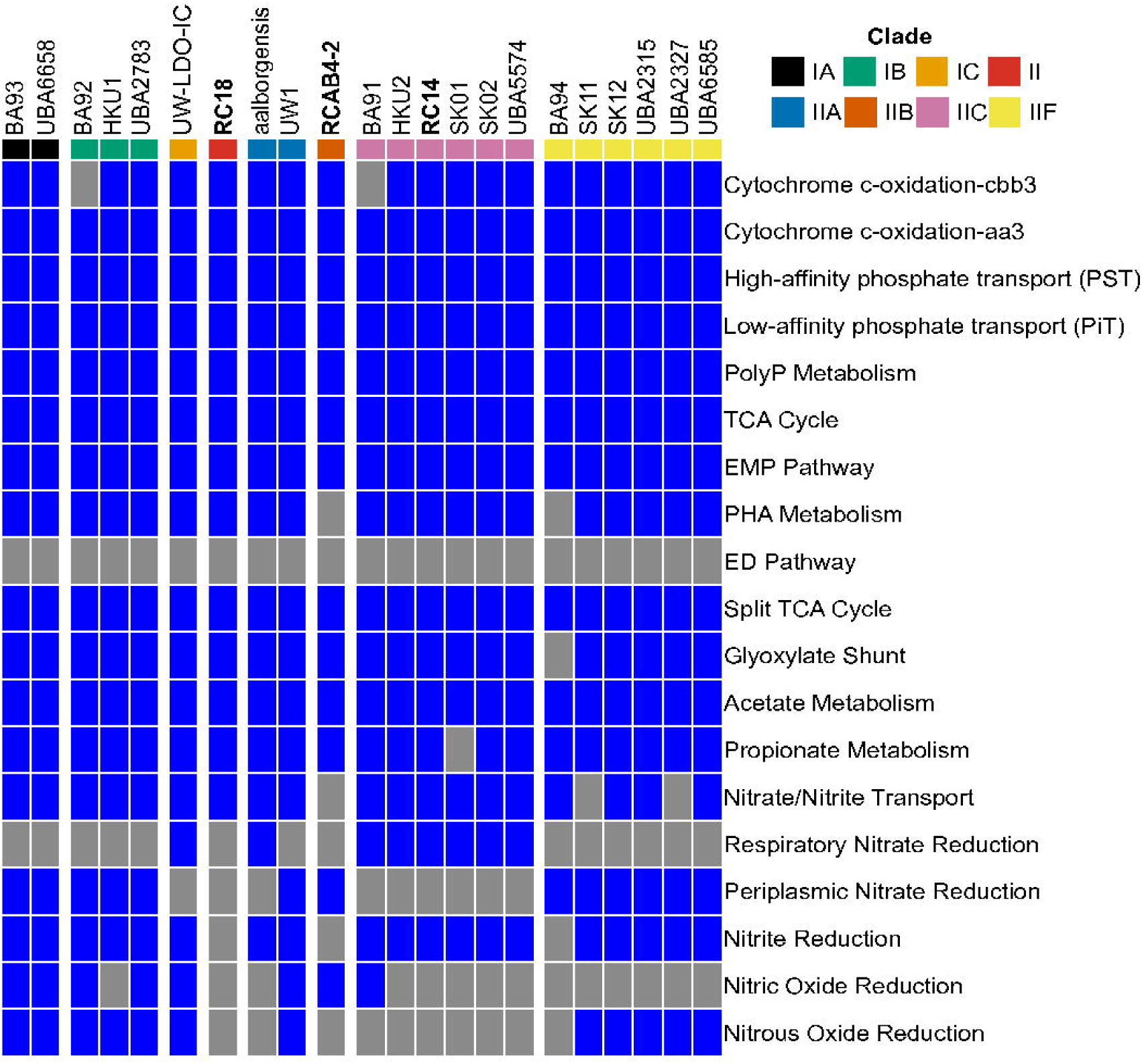
Inventory of pathways in Accumulibacter MAGs associated with EBPR metabolism and denitrification. Blue and grey rectangles represent presence and absence of pathways based on annotation using KEGG.

#### Core EBPR Metabolism

The characteristic metabolism of PAOs in EBPR is the concurrent cycling of carbon and phosphorus through storage and use of intracellular polymers such as glycogen, polyhydroxyalkanoates (PHA) and polyphosphate (polyP). The MAGs extracted in this study (RC14, RC18 and RCAB4-2) were annotated with all genes (Figure 4 & Table S3) considered essential for phosphate transport and polyP metabolism, including *pstSCAB, pitA, ppk1, ppk2* and *ppx*. It has been postulated that the presence of the low-affinity Pit transporter is vital to the PAO-phenotype [65]. The concerted action of these enzymes has been shown to be essential for the PAO phenotype and are present in all Accumulibacter MAGs including the three identified in this study [66, 67]. Genes for PHA synthesis, glycolysis and TCA cycle from both acetyl-CoA and propionyl-CoA were shared among all the MAGs. One of the debated features of the carbon metabolism in Accumulibacter is whether anaerobic glycolysis proceeds via the ED or the EMP pathway [68]. The consistent presence of the EMP pathway and the absence of genes for the ED pathway in the currently available 22 MAGs of Accumulibacter, including 3 MAGs from full-scale facilities in this study, suggests that the EMP pathway is likely the predominant metabolism for anaerobic glycolysis (Figure 4 & Table S3). Another uncertainity in the anaerobic metabolism of EBPR is the source of the reducing equivalents for PHA formation under anaerobic conditions. Current hypothesized pathways include glycolysis, anaerobic operation of the TCA cycle, glyoxylate shunt and split TCA cycle [69–71]. Key enzymes for all these proposed pathways are present in the 3 MAGs from this study (Figure 4). This along with trancriptomic evidence from literature [11, 15, 18, 72, 73] suggests that Accumulibacter likely possess phenotypic versatility and can utilize various pathways in various combinations, depending on the availability of external substrate and intracellular polymers, for their anaerobic metabolism [71, 74, 75].

#### Carbon Metabolism

One of the critical aspects of PAO-type metabolism is the uptake of low molecular weight compounds such as VFAs (primarily acetate and propionate) under anaerobic conditions to form PHA. The 3 Accumulibacter MAGs, obtained in this study, contain the genes necessary for passive transport of acetate and propionate (*actP*) and conversion of acetate and propionate into acetyl-CoA and propionyl-CoA through acetate-coA synthetase (*acsA*) and propionate-coA ligase (*prpE*) respectively (Figure 4 & Table S3).

Full-scale facilities contain a variety of substrates in the wastewater including complex substrates, therefore the ability to use complex substrates can provide a niche for a PAO in full-scale facilities. Most of the MAGs of Accumulibacter, including the 3 MAGs from this study, encode a lactate dehydrogenase gene (*dld*) (except for BA94 and SK01) which catalyzes the conversion of lactate to pyruvate and the reverse reaction. The lactate permease gene (*lctP*/lldP) was identified in aalborgensis, BA91, HKU2, UW1, RC18 and RCAB4-2. The presence of the lactate permease gene, in combination with lactate dehydrogenase, could confer the ability to uptake lactate and convert into pyruvate, acetyl-CoA or propionyl-CoA which can then be used to form PHB/PHV or feed the TCA cycle. Conversion of lactate to pyruvate is able to generate electron equivalents which can also be used for other active transport processes. A recent study [76] showed that Accumulibacter is capable of using lactate to store PHA, however, the authors noticed a deterioration in EBPR performance when feeding lactate presumably associated with a metabolic shift in Accumulibacter from utilizing both polyP and glycogen to utilizing just glycogen (similar to GAOs) since the use of lactate doesn’t require the hydrolysis of polyP to generate ATP.

#### Nitrogen Metabolism

Another trait of interest in Accumulibacter metabolism is the ability to use various oxidized nitrogen species as electron acceptors. Among the available MAGs, only those in Clade IIC MAGs, including MAG RC14 that was dominant in the S2EBPR configuration in this study and one exception from IIA (aalborgensis), contain gene sets for respiratory nitrate reduction (*narGHI*) and nitrite reduction (*nirS*), but did not possess genes for subsequent steps of denitrification. The only MAG that contains genes for a complete denitrification pathway is UW-LDO-IC (Clade IC). Recent work with metatranscriptomics [20] showed that under micro-aerobic conditions and in the presence of nitrate, UW-LDO-IC is able to use both nitrate and oxygen simultaneously as electron acceptors even though the ability to use nitrate as an electron acceptor with exclusive and concurrent uptake of phosphate was not shown.

Most other clades (Clade IIA, IIB, IIF, IA, and IB) contain genes for periplasmic dissimilatory nitrate reduction (*napAB*) system. The Nap system is generally considered to not be coupled with PMF generation and anaerobic respiration. It has been suggested that this system might be involved in aerobic denitrification for roles in adaptation to anaerobic metabolism after transition from aerobic conditions such as in EBPR [78]. There is also evidence that Nap is a dissimilatory enzyme used for optimizing redox balancing [79, 80]. The possession of periplasmic Nap system enzymes in majority of the Accumulibacter clades seem to suggest that these Accumulibacter likely do not use nitrate as an electron acceptor for respiration and denitrification. Rather, the Nap system likely allows CAPs to aerobically reduce nitrate to nitrite in the periplasm, which can be utilized for downstream nitrite reduction or excreted outside the cell.

It is possible that Accumulibacter species with truncated denitrification pathways could use complementary pathways in flanking populations to perform complete denitrification on a community level [77]. Presence of genes for nitric oxide reduction are restricted to a few MAGs in Clade II (UW1, RCAB4-2, BA91) and most Clade I MAGs. In contrast, genes for nitrous oxide reduction are present in almost all Clade IIF, Clade I MAGs and UW1 (Clade IIA). Noticeably, RC18 (present in both the conventional and S2EBPR full-scale systems) did not possess any of the reductases in the denitrification pathway, despite having a completion of 98.2 %.

#### Unique Functions in MAGs from Full-Scale Systems

In this study, we were able to obtain 3 Accumulibacter MAGs from full-scale EBPR communities. It is likely that MAGs from full-scale facilities have some unique functional abilities compared to MAGs from enriched lab-scale bioreactors to enable the organism to survive in the highly dynamic and complex environmental conditions in wastewater treatment systems. We identified genes for dimethyl sulfoxide (DMSO) reduction exclusively in RC18. The pathway included the *dmsA* and *dmsB* subunits of the *dmsABC* gene cluster. The DMSO respiratory pathway has been extensively studied in photo-heterotrophic organisms such as *Rhodobacter* spp. and plays a role in the organism’s redox homeostasis and enables the organisms to utilize a diversity of carbon sources during dark anaerobic growth [80]. Its potential role in Accumulibacter metabolism is unknown and warrants further investigation. We also identified the gene for alkane oxidation, alkane 1-monooxygenase (*alkB*) in RCAB4-2. The production of this enzyme enables the oxidation of alkanes to alcohols making it an important enzyme for bioremediation of petroleum hydrocarbon contaminated environments [81, 82]. Whether this enzyme has a direct impact on EBPR metabolism is unknown.

In this study, for the first time, using integrated amplicon sequencing, oligotyping and genome-resolved metagenomics, we were able to reveal clade-level differences in Accumulibacter communities and associate the differences with two different full-scale EBPR configurations. The results led to the identification and characterization of a distinct and dominant Accumulibacter oligotype - Oligotype 2 (belonging to Clade IIC) and its matching MAG (RC14) associated with S2EBPR configuration. This study demonstrates that oligotyping of the V4 region of the 16S rRNA gene could resolves clade-level differences, congruent with *ppk1*-phylogeny, in Accumulibacter communities in full-scale EBPR systems. The potential advantages of resolving clade-level differences using a high-throughput method such as 16S rRNA gene amplicon sequencing combined with oligotyping is significant since it will enable wider capture, higher-resolution and more cost-effective genotyping of Accumulibacter communities and thus improve our ability to characterize and monitor EBPR processes. Genome-resolved metagenomics also enabled us to retrieve 3 new CAP MAGs from full-scale facilities including two MAGs, RCAB4-2 and RC18, belonging to Clade IIB and to a previously uncharacterized sub-clade of Clade II respectively, that did not have any previous representative genomes. Comparative genomics of the obtained MAGs showed unique functions that could enable niche-partitioning of particular clades under different conditions.

## Acknowledgements

Funding for this research was provided by the Water Environment & Reuse Foundation (Project No: U1R13), Hampton Roads Sanitation District, and Woodard & Curran, Inc. The authors thank Peter Schauer, Adrienne Menniti and Chris Maher (Clean Water Services) for their assistance in this study.

