## Supplementary Data for "Oligotyping and Genome-Resolved Metagenomics Reveal Distinct *Candidatus* Accumulibacter Communities in Full-Scale Side-Stream versus Conventional Enhanced Biological Phosphorus Removal (EBPR) Configurations"

**Running Title:** Oligotyping reveals clade differences of Accumulibacter

Varun N. Srinivasan[1], Guangyu Li[1], Dongqi Wang[1,2], Nicholas B. Tooker[1,3], Zihan Dai[4], Annalisa Onnis-Hayden[1], Ameet Pinto[1], April Z. Gu[1,5*]

[1]Department of Civil and Environmental Engineering, Northeastern University, Boston MA-02115, USA

[2]State Key Laboratory of Eco-hydraulics in Northwest Arid Region, Xi'an University of Technology, Xi'an, Shaanxi 710048, China

[3]Department of Civil and Environmental Engineering, University of Massachusetts-Amherst, Amherst MA-01002, USA

[4]Infrastructure and Environment Division, University of Glasgow, Glasgow G12 8LT, United Kingdom

[5]Civil and Environmental Engineering, Cornell University, Ithaca NY – 14853, USA

[*]**Corresponding Authors**

April Gu, Civil and Environmental Engineering, Cornell University

**SUPPLEMENTARY DATA**

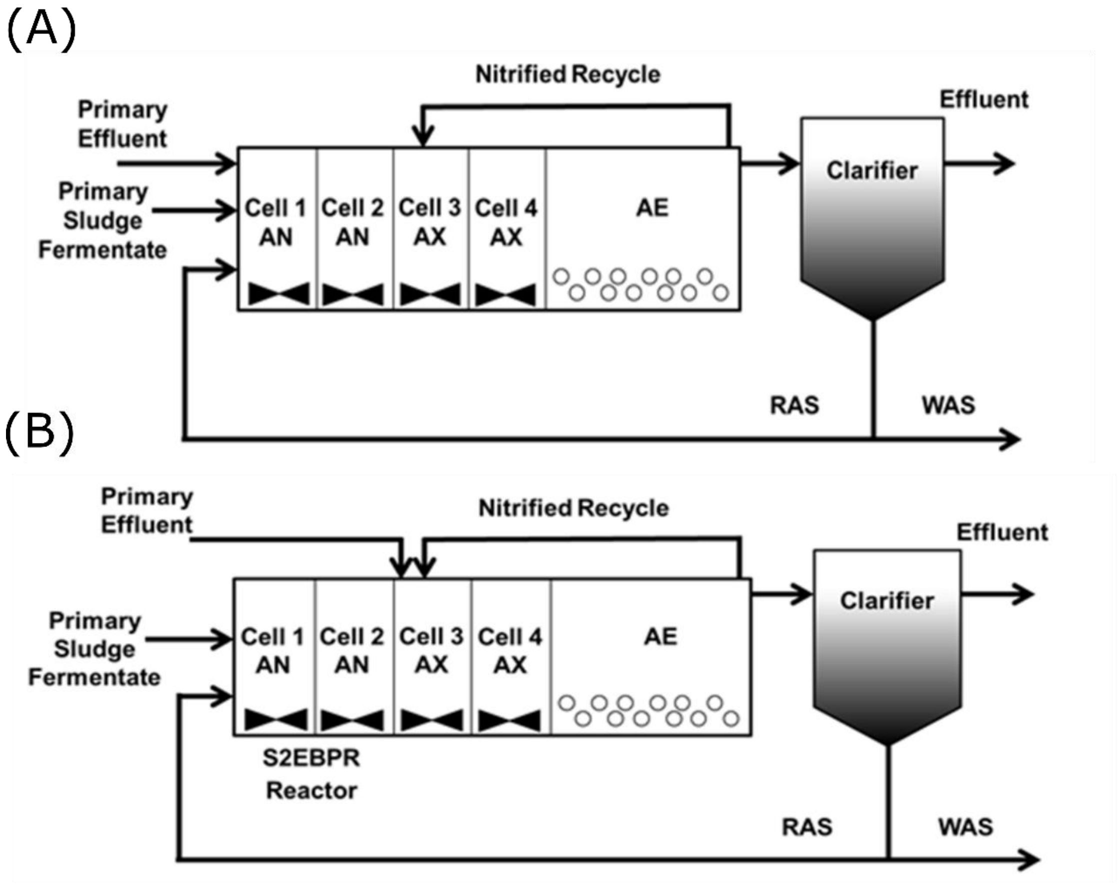


**Figure S 1: Flow schematic of (A) conventional A2O (B) S2EBPR SSRC configurations.**

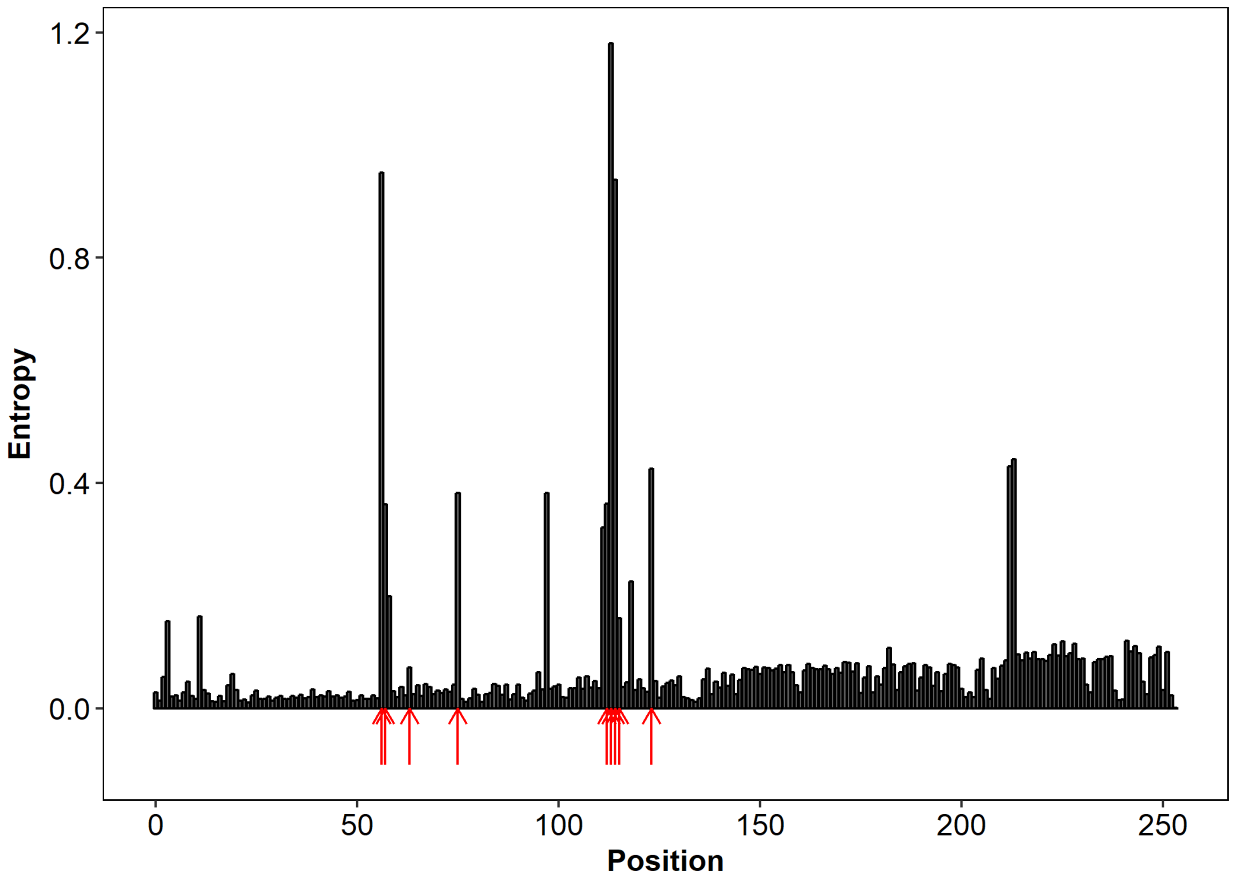


**Figure S 2: Results of entropy analysis using the oligotyping pipeline according Eren et al. [5] with Accumulibacter sequences targeting the V4 region of the 16S rRNA gene. Arrows indicate nucleotide positions used for oligotyping.**

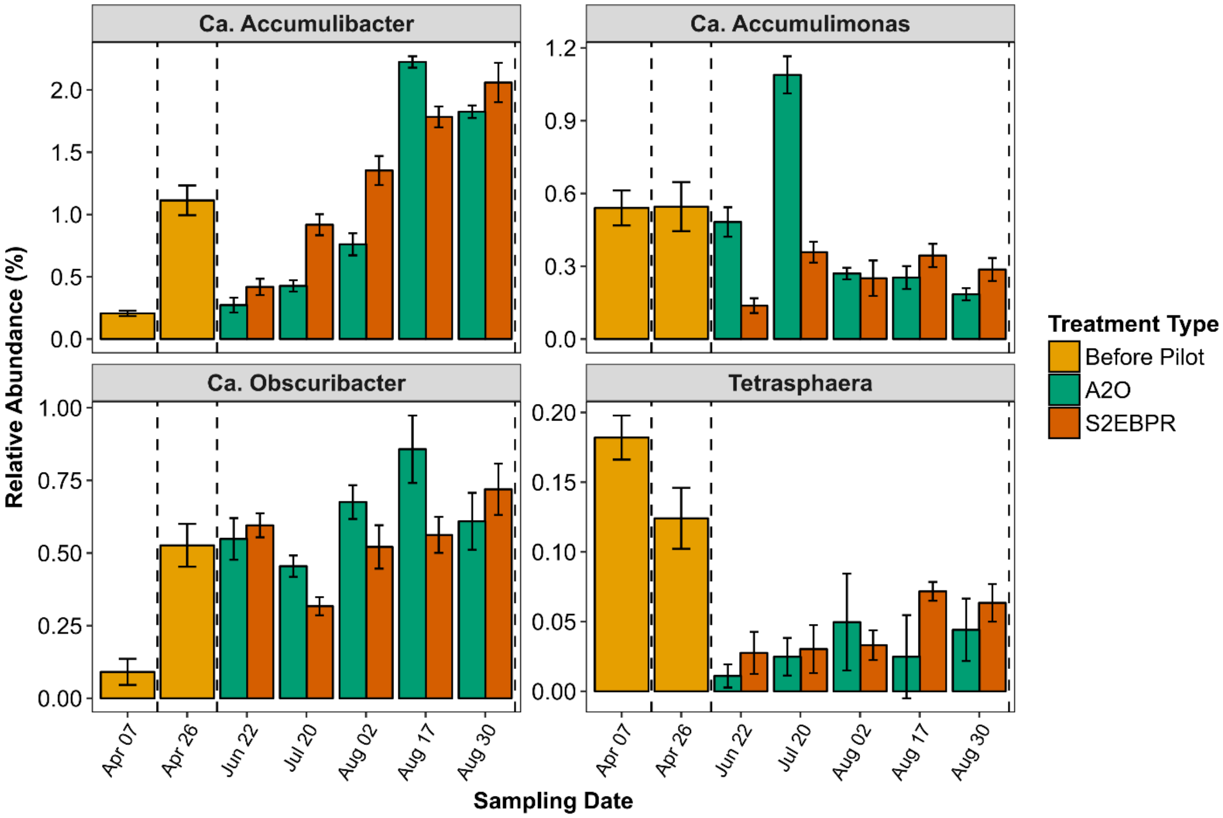


**Figure S 3: Relative abundance of putative known PAOs (based on the MiDAS database) before the pilot and during the pilot.**

**Table S 1: Oligotype sequences and corresponding names used in figures.**

| Oligotype | Name |
|---|---|
| GGTTCCAGT | Oligotype 1 |
| GGTTCTAGT | Oligotype 2 |
| GTTTACGGT | Oligotype 3 |
| TGTGCAAGA | Oligotype 4 |
| TGTTCAAGT | Oligotype 5 |
| TGTTCAGGT | Oligotype 6 |
| TTTTAAAGT | Oligotype 7 |
| TTTTAAGAT | Oligotype 8 |
| TTTTAAGGG | Oligotype 9 |

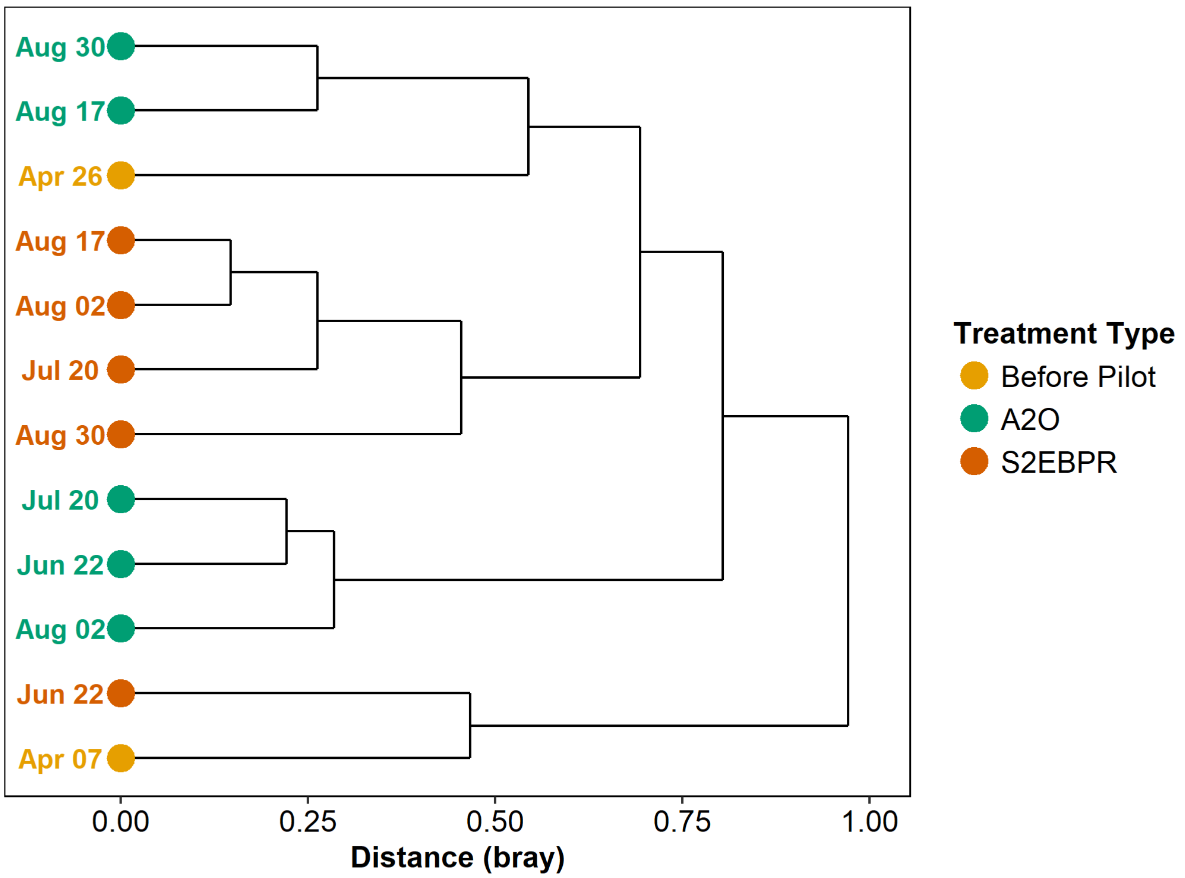


**Figure S 4: Hierarchical Clustering of Samples using the Bray Curtis Distance Matrix of Oligotype Counts**

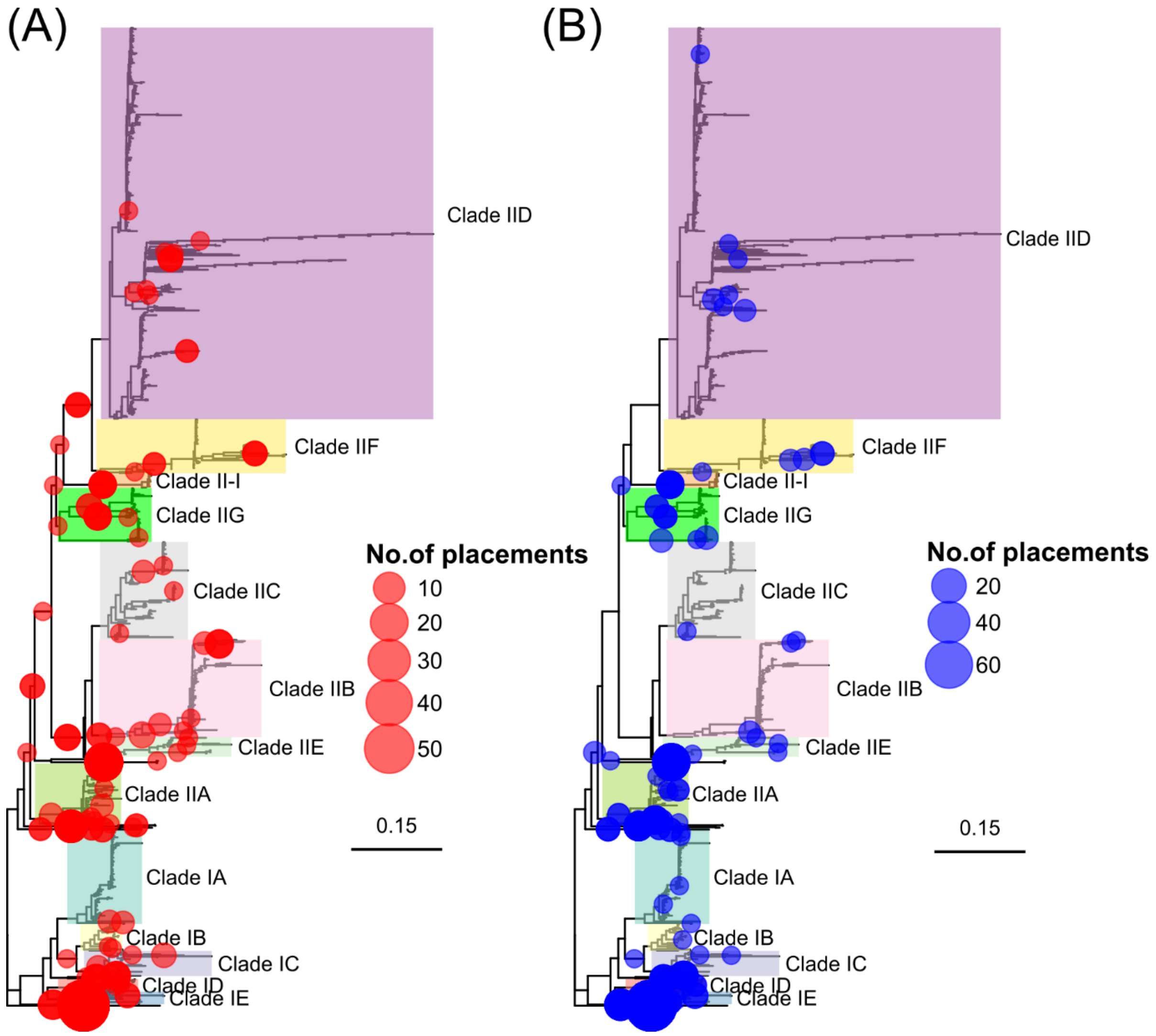


**Figure S 5: (A) Placements of *ppk1* gene sequences extracted from the A2O assembly (B) Placements of *ppk1* gene sequences extracted from the S2EBPR assembly**

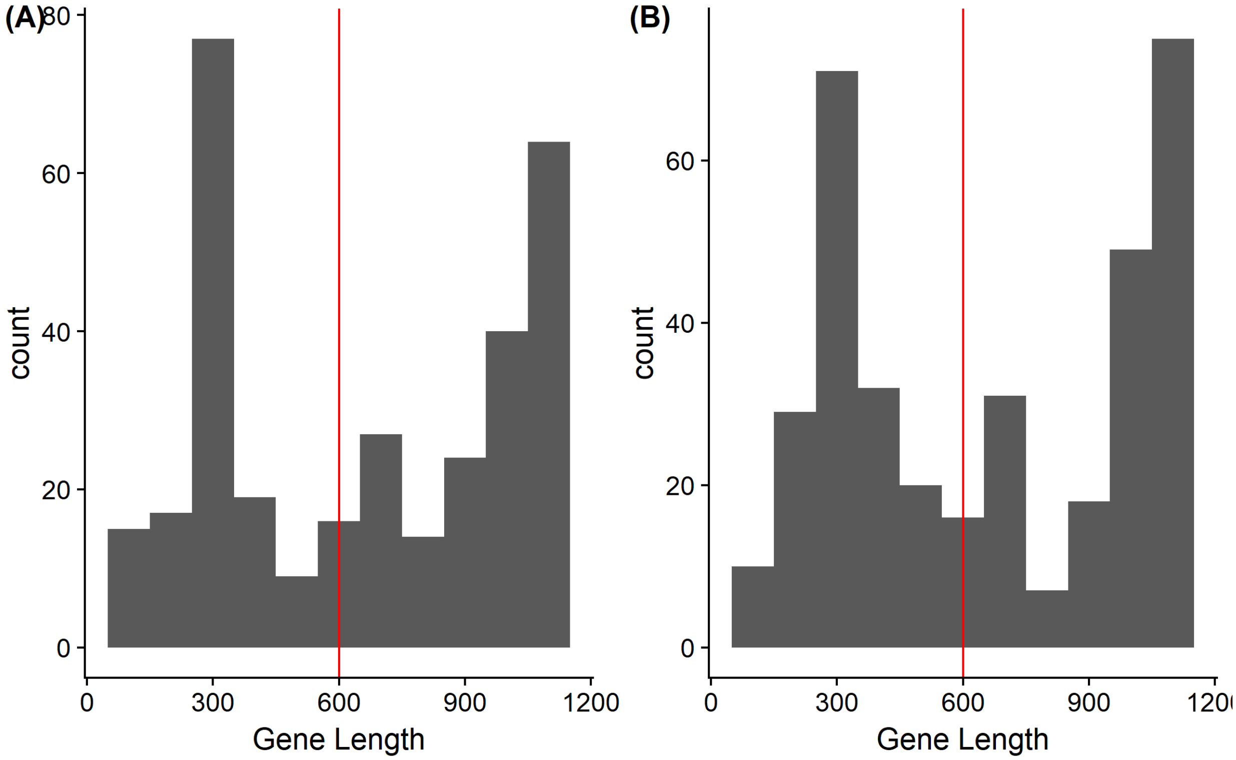


**Figure S 6: Histogram of *ppk1* gene sequence lengths extracted from (A) A2O assembly (B) S2EBPR Assembly. The red line indicates the minimum length (600 bp) of *ppk1* gene sequences used for analysis.**

**Table S 2: Genome Characteristics for MAGs from This Study.**

| MAG | Completeness (%) | Redundancy (%) | Size (Mbp) | # of contigs | N50 (Kbp) |
|---|---|---|---|---|---|
| RC14 | 99 | 0.2 | 4.57 | 197 | 39 |
| RC18 | 98.2 | 0.0 | 3.43 | 127 | 38.7 |
| RCAB4-2 | 77.2 | 3.1 | 3.03 | 608 | 5.3 |

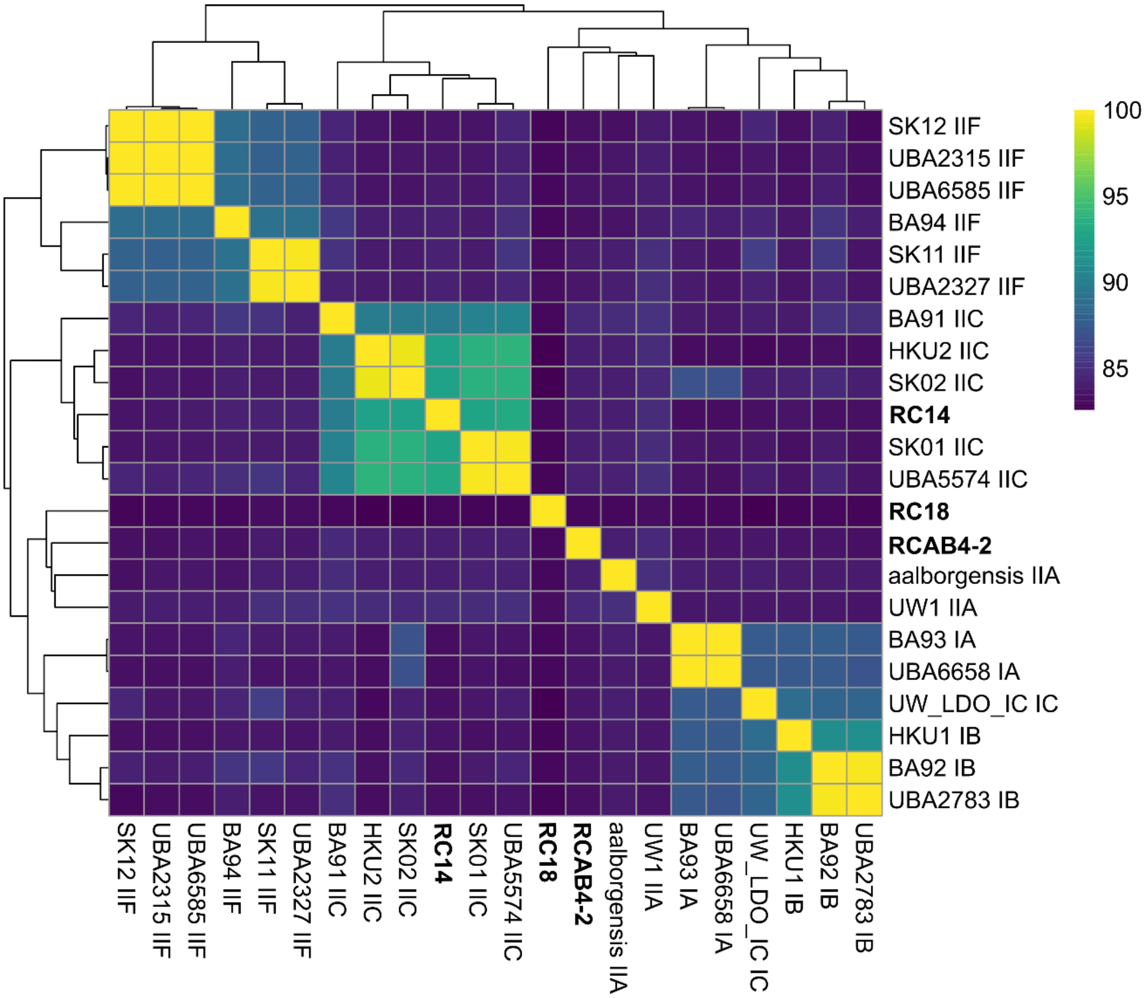


**Figure S 7: Heatmap of Average Nucleotide Identity (ANI) of all Available Accumulibacter MAGs. The full-scale MAGs from this study are highlighted in bold.**
