## Supplementary Methods for "Oligotyping and Genome-Resolved Metagenomics Reveal Distinct *Candidatus* Accumulibacter Communities in Full-Scale Side-Stream versus Conventional Enhanced Biological Phosphorus Removal (EBPR) Configurations"

**Running Title:** Oligotyping reveals clade differences of Accumulibacter

Varun N. Srinivasan[1], Guangyu Li[1], Dongqi Wang[1,2], Nicholas B. Tooker[1,3], Zihan Dai[4], Annalisa Onnis-Hayden[1], Ameet Pinto[1], April Z. Gu[1,5*]

[1]Department of Civil and Environmental Engineering, Northeastern University, Boston MA-02115, USA

[2]State Key Laboratory of Eco-hydraulics in Northwest Arid Region, Xi'an University of Technology, Xi'an, Shaanxi 710048, China

[3]Department of Civil and Environmental Engineering, University of Massachusetts-Amherst, Amherst MA-01002, USA

[4]Infrastructure and Environment Division, University of Glasgow, Glasgow G12 8LT, United Kingdom

[5]Civil and Environmental Engineering, Cornell University, Ithaca NY – 14853, USA

**[*]Corresponding Authors**

April Gu, Civil and Environmental Engineering, Cornell University

### SUPPLEMENTARY METHODS

#### *Rock Creek Pilot*

Full-scale pilot testing was conducted during summer 2016 at the Rock Creek Facility (Hillsboro, Oregon). Two parallel trains with separated secondary clarifiers were operated with the same influent wastewater with side-by-side operation of conventional A2O and S2EBPR SSRC configurations (Figure S1). Before pilot testing was started, both the trains were operated in the A/O configuration and the RAS from the two trains were blended together. At the start of pilot testing, the RAS from the two treatment trains were separated and the trains were operated independently. The S2EBPR configuration implemented was side-stream RAS fermentation with supplemental carbon addition (SSRC). In this configuration, 100% of the RAS was diverted to the side-stream reactor (Figure S1) for a total anaerobic HRT of 1.5 hours with primary sludge fermentate addition. The primary effluent was fed into the anoxic zone of the main-stream which was followed by an aerobic zone. Initially, the anaerobic side-stream reactor was completely mixed. After 2 months (April 26th – June 21st), the side-stream reactor was switched to intermittent mixing once per week for 10 minutes. The conventional EBPR configuration was an A2O process with an anaerobic HRT of 0.7 hours. Throughout pilot testing, process performance parameters were routinely monitored. More detailed information about design, operation and process performance is available in previously published literature [1–3].

#### *Amplicon Sequencing Data Analysis*

The raw Fastq files were cleaned using Sickle 1.33 [4] with a minimum window quality score of 20. The sequences were trimmed to remove primers and barcodes, quality filtered using sickle v1.33 with a minimum quality score of 20 and analyzed as described in Kozich et al., [5]. Briefly, the quality-filtered sequences were assembled in mothur [6] and aligned to SILVA v123 database. The alignment was screened to remove poorly aligned sequences using vertical = T and trump = . options in mothur. Chimeras were remove using the UCHIME [7] algorithm available through mothur and clustered into OTUs at sequence similarity cutoff of 97% using the OptiClust method. The sequences were classified using the Naive Bayesian Classifier (80% confidence threshold) using the Silva database (v123) and consensus taxonomy of OTUs was determined using the 80% cutoff using the MiDAS (v123) database.

The relative abundance of each OTU was determined by summing the total number of OTU associated reads and dividing by the total number of reads in the sample. To identify putative PAOs and GAOs (defined as organisms with glycogen-accumulating capabiity and no known phosphate-accumulating capability), the MiDAS field guide website (http://midasfieldguide.org/en/search/) was used. The following filters were used to download a list of organisms with PAO-like function (PAO-positive) and GAO-like function (GAO-positive and PAO-negative). Diversity indices were calculated using the vegan package [8] in R.

***Reconstruction of 16S rRNA Gene Sequences and Oligotype Sequences***

EMIRGE [9] was used to reconstruct 16S rRNA gene sequences from the paired-end metagenomic data. Mean insert sizes and standard deviations were determined using Picard (CollectInsertSizeMetrics) and used in EMIRGE with 40 iterations and with the flag –j 1 (to get unique sequences). The sequences were then renamed using emirge_rename_fasta.py script and concatenated for dereplication and classification. A total of 489 sequences were obtained from all 6 samples. The sequences were first screened using the screen.seqs command in mothur to remove sequences shorter than 1000 bp , longer than 1700 bp and with any ambiguous base pairs. The sequences were then dereplicated in mothur using unique.seqs  command. A total of 394 unique sequences were left. The consensus taxonomy for these sequences was determined using the Naïve Bayesian Classifier in mothur with 80% cutoff using the MiDAS (v123) database. The sequences classified as Ca. Accumulibacter were extracted, an in-silico PCR was performed using the 515F and 806R primers and a nucleotide blast with 100 % identity criteria against the oligotype sequences was performed to extract exact sequence-matches to the oligotypes. Exact sequence matches were then aligned with the reference sequences from He et al., [10]. The alignment was trimmed using trimAl [11] with the –gappyout flag which resulted in an alignment length of 1476 columns. A phylogenetic tree was constructed in MrBayes 3.2 [12] with a General Time Reversible Model with a gamma-distributed rate variation with 3,000,000 generations and a sampling frequency of 100. A consensus tree was generated after discarding all trees generated before burn-in set in.

12. Ronquist F, Huelsenbeck JP. MrBayes 3: Bayesian phylogenetic inference under mixed

models. *Bioinformatics* 2003; **19**: 1572–1574.
